# Norovirus replication in human intestinal epithelial cells is restricted by the interferon-induced JAK/STAT signalling pathway *and* RNA Polymerase II mediated transcriptional responses

**DOI:** 10.1101/731802

**Authors:** Myra Hosmillo, Yasmin Chaudhry, Komal Nayak, Frederic Sorgeloos, Bon-Kyoung Koo, Alessandra Merenda, Reidun Lillestol, Lydia Drumright, Matthias Zilbauer, Ian Goodfellow

## Abstract

Human noroviruses (HuNoV) are a leading cause of viral gastroenteritis worldwide and a significant cause of morbidity and mortality in all age groups. The recent finding that HuNoV can be propagated in B cells and mucosa derived intestinal epithelial organoids (IEOs), has transformed our capability to dissect the life cycle of noroviruses. Using RNA-Seq of HuNoV infected intestinal epithelial cells (IECs), we have found that replication of HuNoV in IECs results in interferon-induced transcriptional responses and that HuNoV replication in IECs is sensitive to IFN. This contrasts with previous studies that suggest that the innate immune response may play no role in the restriction of HuNoV replication in immortalised cells. We demonstrate that the inhibition of JAK1/JAK2 enhances HuNoV replication in IECs. Surprisingly, targeted inhibition of cellular RNA polymerase II-mediated transcription was not detrimental to HuNoV replication, but enhanced replication to a greater degree compared to blocking of JAK signalling directly. Furthermore, we demonstrate for the first time that IECs generated from genetically modified intestinal organoids, engineered to be deficient in the interferon response, are more permissive to HuNoV infection. Together our work identifies the IFN-induced transcriptional responses restrict HuNoV replication in IECs and demonstrates that the inhibition of these responses by modifications to the culture conditions can greatly enhance the robustness of the norovirus culture system.

**Importance:** Noroviruses are a major cause of gastroenteritis worldwide yet the challenges associated with their growth culture has greatly hampered the development of therapeutic approaches and has limited our understanding of cellular pathways that control infection. Here we show that human intestinal epithelial cells, the first point of entry of human noroviruses into the host, limit virus replication by the induction of the innate responses. Furthermore we show that modulating the ability of intestinal epithelial cells to induce transcriptional responses to HuNoV infection can significantly enhance human norovirus replication in culture. Collectively our findings provide new insights into the biological pathways that control norovirus infection but also identify mechanisms to enhance the robustness of norovirus culture.

## Introduction

The induction of the host innate response plays an essential role in the suppression of pathogen infection. The synthesis of interferons (IFN) and the subsequent signalling cascades that leads to the induction of IFN-stimulated genes (ISGs), determine the outcome of viral infection (1, 2). An understanding of the mechanisms underlying the interplay between pathogens and innate immune responses is vital to understanding viral pathogenesis and can greatly aid the identification of potential therapeutic and/or preventive strategies.

Human noroviruses (HuNoV) are widely recognised as the leading cause of viral gastroenteritis worldwide (3). Noroviruses are classified into at least seven genogroups based on the sequence of the major capsid protein VP1 and regions within ORF1 (3–5). HuNoVs belong to one of three norovirus genogroups (GI, GII, or GIV), which are further divided into >25 genetic clusters or genotypes (6–8). Epidemiological studies reveal that over 75% of confirmed human norovirus infections are associated with HuNoV GII (9, 10). Whilst norovirus gastroenteritis typically results in an acute and self-limiting disease, the socioeconomic impact in both developed and developing countries is estimated to be more than $60.3 billion per annum (11). HuNoV infection is particularly severe and prolonged in immunocompromised patients, including young children, elderly, or patients receiving treatment for cancer. In these cases infections can last from months to years (12, 13).

Our understanding of the molecular mechanisms that control HuNoV infection has been limited by the lack of robust culture systems that facilitate the detailed analysis of the viral life cycle. As a result, murine norovirus (MNV) and other members of the *Caliciviridae* family of positive sense RNA viruses, such as feline calicivirus (FCV) and porcine sapovirus (PSaV), are often used as surrogate models (14–17). MNV, FCV and PSaV can all be efficiently cultured in immortalised cells and are amenable to reverse genetics (16–20). These model systems have been critical to understanding many aspects of the life cycle of members of the *Caliciviridae* (15).

Recent efforts have led to the establishment of two HuNoV culture systems based on immortalised B cells (21, 22) and intestinal epithelial cells (IECs) generated from biopsy-derived human intestinal epithelial organoids (IEOs) (23). Whilst authentic replication of HuNoV can be observed in both the B-cell and IEC-based culture systems, repeated long-term passage of HuNoV and the generation of high titre viral stocks is not possible, suggesting that replication is restricted in some manner. In the current study we sought to better understand the cellular response to HuNoV infection and to identify pathways that restrict HuNoV replication in organoid-derived IECs. Using RNA-Seq we observed that HuNoV infection of IECs results in an interferon-mediated antiviral transcriptional response. We show for the first time that HuNoV replication in IECs is sensitive to both Type I and III interferon and that HuNoV replication is restricted by virus-induced innate response. Pharmacological inhibition of the interferon response or genetic modification of organoids to prevent the activation of the interferon response significantly improved HuNoV replication in IECs. Furthermore, we show that ongoing HuNoV replication is enhanced by the inhibition of RNA Pol II mediated transcription. Overall this work provides new insights into the cellular responses to HuNoV infection of the gut epithelium and identifies modifications to the HuNoV culture system that significantly enhances its utility.

## Materials and Methods

### Stool samples

Stool specimens were anonymized with written consent from patients at Addenbrooke’s Hospital, Cambridge, who tested positive of HuNoV infection. Stool samples were diluted 1:10 (wt/vol) with phosphate buffered saline (PBS) and processed as described (23). Briefly, 10% stool suspensions were vigorously vortexed for 1 min and sonicated three times for 1 min (50/60 Hz, 80W). Homogenous fecal suspensions were centrifuged at 1,500xg for 10 min at 4 C. The supernatants were serially passed through 5 μm, 1.2 μm, 0.8 μm, 0.45 μm and 0.22 μm filters (Millex-GV syringe filter units). Stool filtrates were aliquoted and stored at -80 C until used.

### Human intestinal organoids

Following ethical approval (REC-12/EE/0482) and informed consent, biopsies were collected from the proximal duodenum (D) or terminal ileum (TI) from patients undergoing routine endoscopy. All patients included had macroscopically and histologically normal mucosa. Biopsy samples were processed immediately and intestinal epithelial organoids generated from isolated crypts following an established protocol as described previously (23–25). Intestinal organoids were grown in proliferation media (Table S1) as described (24). Organoids were typically grown for 7-9 d prior to passage at ratios of 1:2 to 1:3.

Following the establishment of organoid cultures, differentiated IEC monolayers were generated on collagen-coated wells in differentiation media (Table S1) as described (23). Following 5 days of differentiation, confluent monolayers of differentiated IECs were infected. Differentiation was assessed by RT-qPCR at various time post infection by assessing the levels of the stem cell marker LGR5, a mature enterocyte marker alkaline phosphatase (ALP) and epithelial cell marker villin (VIL). Data were normalised to the housekeeping gene hypoxanthine phosphoribosyltransferase 1 (HPRT1).

### Cell lines and reagents

L-WNT 3A expressing cell lines were used to produce WNT conditioned media as a component of the proliferation media and were propagated in low glucose DMEM (Life technologies), 10% fetal calf serum (FCS), 1% penicillin-streptomycin (P/S) and Zeocin (125 μg/ml) at 37°C with 5% CO_2_. WNT-conditioned media was collected from cells grown in the absence of Zeocin. The activity of WNT3a in conditioned media was assessed using a luciferase reporter assay reliant on a Wnt3A responsive promoter (HEK 293 STF, ATCC CRL-3249).

293T-RSPO-V5 cells were used to produce the R-Spondin 1 (RSPO1) conditioned media. 293T-RSPO-V5 cells were propagated in DMEM (Life technologies), 10% fetal calf serum (FCS), 1% penicillin-streptomycin (P/S) and Zeocin (300 μg/ml) at 37°C with 5% CO_2_. RSPO1-conditioned media were collected from passage of cells in conditioned media containing DMEM/F12 (Life technologies),1% penicillin-streptomycin (P/S), 10mM HEPES (Life technologies), and 1x glutamax (Life technologies).

Components of the proliferation and differentiation media are described in Table S1. The commercial sources of Interferons (IFNαA/D, Sigma; IFNβ1, IFNλ1, and IFNλ2, Peprotech) and IFN inhibitors Ruxolitinib (Invivogen) and Triptolide (Invivogen) are detailed in Table S2.

### Lentivirus vector particle production and transduction

Lentivirus transfer vectors encoding the BVDV NPro and PIV5 V proteins were a gift from Professor Steve Goodbourn (St. George’s Hospital, University of London). The transfer vectors were used to generate vesicular stomatitis virus G-protein-pseudotyped lentiviral particles by transfection of 293T cells with psPAX2 and pMD2.G helper plasmids. Human IEOs were then transduced with lentivirus-containing supernatants following published protocols (18). Transduced organoids were selected with puromycin (2 μg/ml) and organoid clones were selected by limiting dilution and subsequent functional analysis.

### HuNoV infection

Differentiated monolayers in 48-well plates were infected in biological duplicates or triplicate as described in the text. HuNoV stool filtrates containing ∼1 x 10^6^ viral RNA copies, determined by RT-qPCR, were added to each well and incubated 37 °C for 2 h, prior to being washed twice with serum-free media and overlaid with 250 µL of differentiation media containing 200 µM GCDCA. Where required, wells were supplemented with either DMSO, IFN or pharmacological inhibitors as described in the text. Samples were typically harvested at 48 hours post infection for analysis.

Inactivated HuNoV-containing stool filtrates were prepared placing stool filtrate into multiple 24-well plates to a fluid depth of 10 mm and exposing to 4000 mJ from a UV source for 12 min at 4 °C. Loss of viral infectivity was confirmed by infection of monolayers and by comparison of viral titres observed after 48 hours post infection with that obtained using the well-characterised RNA polymerase inhibitor 2-CMC (26).

### qRT-PCR and qPCR analysis

Gene-specific primers and probes against the cellular mRNAs HPRT, LGR5, ALP, and VIL (Thermo Fisher Scientific) were used to evaluate differentiation by RT-qPCR. Samples were analysed by technical duplicate qPCR reactions and the results averaged.

HuNoV GII specific primers previously reported (27) were used in Taqman-based qRT-PCR assay to detect HuNoV replication in organoid cultures. The levels of HuNoV mRNA were determined based on absolute quantitation against a standard curve generated using *in vitro* transcribed RNA from a full length cDNA clone of a GII.4 HuNoV. Each individual biological sample was analysed by qRT-PCR in technical duplicate alongside additional no template negative controls. Data were collected using a ViiA 7 Real-Time PCR System (Applied Biosystems).

### RNA library preparation and sequencing

Total cellular RNA was extracted from IECs using Trizol (Invitrogen) and genomic DNA was removed by DNase I digestion (TURBO DNA-free™ Kit, Ambion, AM1907). RNA integrity was assessed via an Agilent 2200 TapeStation system using RNA ScreenTape reagents (Agilent Technologies, catalog number: 5067-5576/77). Libraries were prepared for sequencing by Cambridge Genomics Services, 300 ng of total RNA using a TruSeq stranded mRNA kit (Illumina Technologies, catalog number: 20020595). Libraries were quantified by qPCR, pooled and sequenced with 75 basepair single reads to a depth ranging from 13 to 55 million reads per sample on a Illumina NextSeq 500 using a High Output 75 cycles kit (Illumina, catalog number: FC-404-2005).

### Data analysis

Raw reads were inspected with FastQC. Adapters and low quality sequences were removed using Trimmomatic version 0.33 using the following parameters:ILLUMINACLIP:TruSeq3-SE:2:30:10LEADING:3TRAILING:3SLIDIN GWINDOW:4:15 MINLEN:36. Transcript level quantification for each sample was obtained using the kallisto software (28) against the human transcriptome GRCh38.p12 (Ensembl release v92; accessed 22/03/2018) and genes with average transcripts per million less than 1 in both control and virus infection conditions were excluded from downstream analysis. Read counts were normalized using the trimmed mean of M-values normalization method (29) and log_2_ count per million (CPM) were obtained using calcNormFactors and voom (30) functions of edgeR (29) and limma (31) packages, respectively. Student’s t-tests were then applied for each transcript. Finally, p-values were adjusted for false discoveries due to simultaneous hypotheses testing by applying the Benjamini–Hochberg procedure (FDR) (32). Transcripts with a FDR lower than 0.01 and log2FC greater than 1 (FC=2) were considered as differentially expressed. Heatmap of gene expression for significant genes across comparisons was generated using the R package pheatmap version 1.0.10. Levels of expression change are represented by a colour gradient ranging from blue (low increase in gene expression) to red (high increase in gene expression). Gene ontology term and Reactome pathway enrichment analyses were performed with the clusterProfiler R package version 3.8.1 (33) and represented using the R package pheatmap as above.

The RNA-Seq data obtained in this study have been deposited the Gene Expression Omnibus (GEO, http://www.ncbi.nlm.nih.gov/geo) with the accession number GSE117911. <u>Reviewer access can be obtained by entering the token“mholyaqyrrurvof”.</u> In addition, all sequence reads were deposited in the NCBI Sequence Read Archive Database (SRA, http://www.ncbi.nlm.nih.gov/sra) and are associated with the accession number PRJNA483555.

### Statistical analysis and software

Statistical analyses were performed on triplicate experiments using the two-tailed Student t-test (Prism 6 version 6.04). Figures were generated using Inkscape and Prism 8 version 8.0.2.

## Results

### Human norovirus replicates productively in differentiated intestinal epithelial cells from the human proximal and distal small bowel

Building on previous studies reporting the replication of HuNoV in IECs, we set out to better understand the cellular response to HuNoV infection and to identify pathways that restrict HuNoV replication in IECs. We established IEO cultures using mucosal biopsies obtained from several gut segments of the small intestine, the proximal duodenum and terminal ileum. Given the importance of fucosyltransferase expression on HuNoV susceptibility (34–36), lines were established from FUT2 positive individuals. Intestinal crypt cells were isolated and used to generate small IEOs (Fig. 1A). The 3D-organoid structures are allowed to self-organize and differentiate within Matrigel using optimized proliferation medium as described (Table S1) (24). The established organoid lines were typically cultured for 7-9 d and expanded at passage ratios of 1:2 or 1:3. As expected, during the first three days of culture the intestinal organoids initially formed small cystic structures with a central lumen, lined with epithelial cells (Fig. 1A). By day 5 more convoluted structures formed, the nature of which varied from line to line (Fig. 1A).

**Figure 1:**
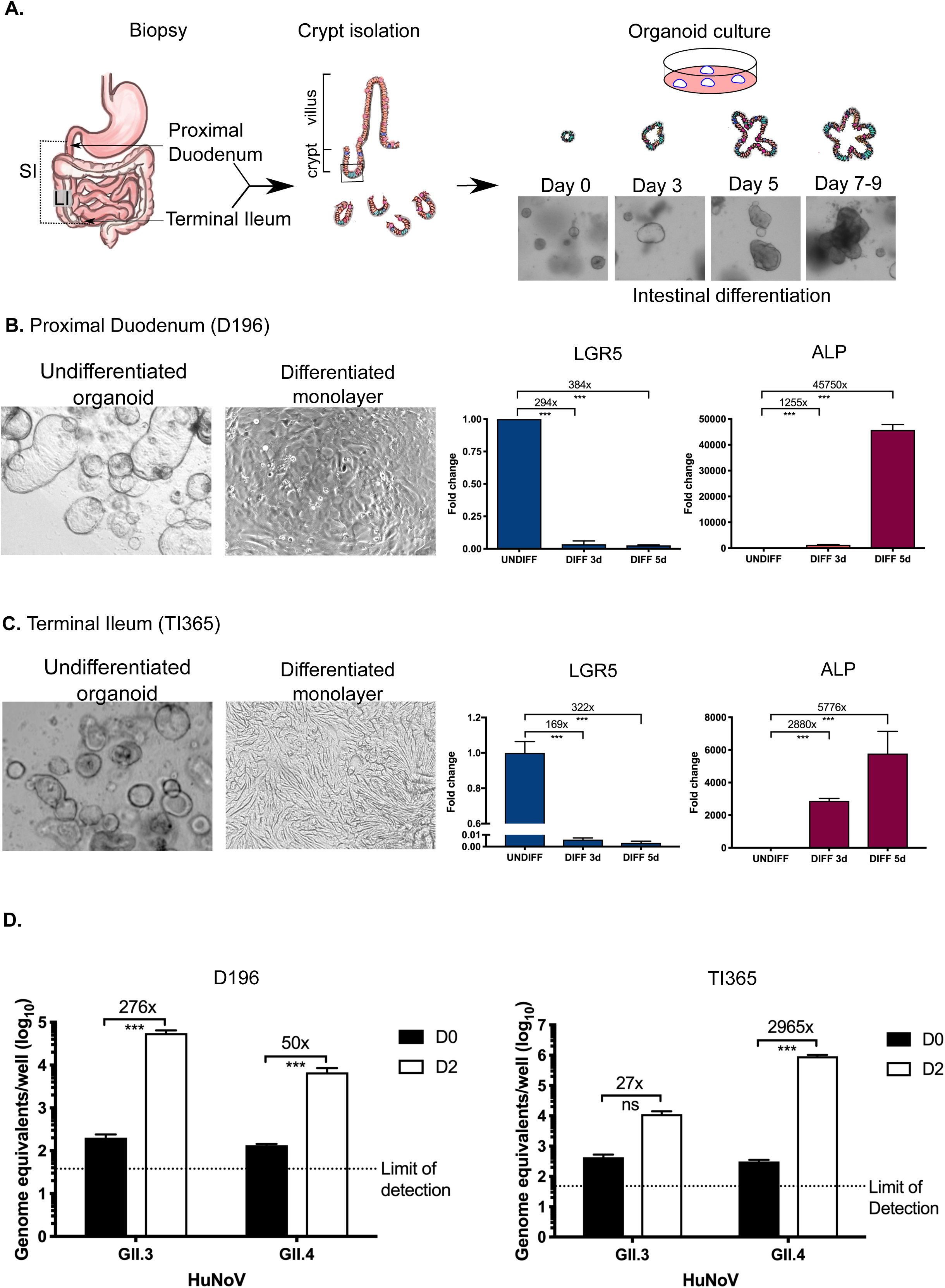
Overview of the human norovirus culture system. A) Schematic of the intestinal crypt isolation procedure leading to the production of intestinal organoids. Following isolation by biopsy, crypts were plated into Matrigel as described in the text and imaged by light microscopy. B, C) differentiation of intestinal organoids from the duodenum and terminal ileum into intestinal epithelial cells (IEC) monolayers is accompanied by loss of the stem cell marker LGR5 and increased intestinal alkaline phosphatase (AP) expression. Intestinal organoid lines were plated onto collagen coated plates as described in the text and the relative levels of LGR5 or ALP quantified by RT-qPCR. Expression levels are shown relative to the undifferentiated cells extracted on day 0 of plating. D) Infection of differentiated IECs from the duodenum (D196) and terminal ileum (TI365) with two clinical isolates of human norovirus (GII.3 and GII.4). Infection was assessed by the quantification of absolute viral RNA levels by RT-qPCR and is shown as both absolute values (D) and fold the increase in viral RNA levels when comparing day 0 to day 2 (D2).

To assess the replication efficiency of HuNoV replication in differentiated IECs, 7-9 day-old organoids were plated onto collagen-coated plates, then Wnt and RSpo removed to drive differentiation. To examine the degree of differentiation and to confirm the presence of enterocytes in the monolayers, we examined the mRNA levels of LGR5 and ALP in IEC monolayers generated from both duodenum and ileum (Fig. 1B and 1C). As shown in Fig. 1B, the levels LGR5 mRNA in proximal duodenum decreased by ∼294- to 384-fold, whereas ALP mRNA increased by 1200- to 45000-fold following the removal of Wnt and RSpo. Similarly, the differentiation of IECs derived from the terminal ileum was confirmed by an increased in ALP mRNA and a concomitant decreased in LGR5 (Fig. 1C). These results confirmed that the IEC monolayers had undergone differentiation and confirmed the presence of enterocytes in the differentiated monolayers.

To assess HuNoV replication in the human IEC monolayers, filtered stool samples containing genogroup II HuNoV strains were inoculated onto differentiated monolayers generated from either duodenum and terminal ileum-derived intestinal organoids. Following a 2 hour adsorption period, the inoculum was removed by washing and the monolayers maintained in differentiation media with the bile acid GCDCA for 2d. While previous observations indicated that some strains of HuNoV do not require bile acids for infection, GCDCA was included to maintain a physiologically relevant environment and to control for any effect of bile acids on gene expression. Replication of HuNoV was then assessed by comparing viral RNA levels present in cultures at 2 h post infection (Day 0, D0) to 48 hours post infection (Day 2, D2). In duodenal IEC monolayers, viral RNA levels of both GII.3 and GII.4 HuNoV strains increased by ∼1.5 to 2 log_10_ over the 2 day period (Fig. 1D). Similar levels of viral replication were observed in IEC monolayers derived from terminal ileum organoids, resulting in ∼1.3 to 3.5 log_10_ increases in viral RNA levels (Fig. 1D).

### Norovirus infection of human intestinal epithelial cells induces the innate immune response

The development of a stem-cell derived culture system for HuNoV provides the first opportunity to characterize the cellular pathways that restrict norovirus replication at their primary site of entry into the host, namely the gut epithelium (23). Whilst a previous report suggested that replication of HuNoV did not induce a robust interferon response in immortalized cells (37), inefficient replication and mutations commonly found in cell lines that compromise their ability to respond to viral infection, may have confounded this observation. Whilst there are limited examples in the literature, there is evidence that natural HuNoV infection results in the production of pro- and anti-inflammatory cytokines (38). We have also recently reported that HuNoV replication in Zebrafish also resulted in a measurable innate response (39).

To examine the effect of HuNoV replication on the stimulation of the IFN-induced innate response, we initially assessed the mRNA levels of two candidate interferon stimulated genes (ISGs), human viperin and ISG 15. Viperin and ISG15 mRNAs increased significantly following HuNoV infection (Fig. 2A). To confirm that induction was specifically caused by active HuNoV replication, UV-inactivated HuNoV stool filtrates were used alongside to control for non-specific effects. In contrast to live virus inoculated IEC monolayers, induction of viperin and ISG15 was not seen in cells infected with UV-inactivated virus, confirming this induction as virus-specific (Fig.2A). These results indicate for the first time that active replication of HuNoV in IECs readily stimulates the interferon-induced innate response.

**Figure 2:**
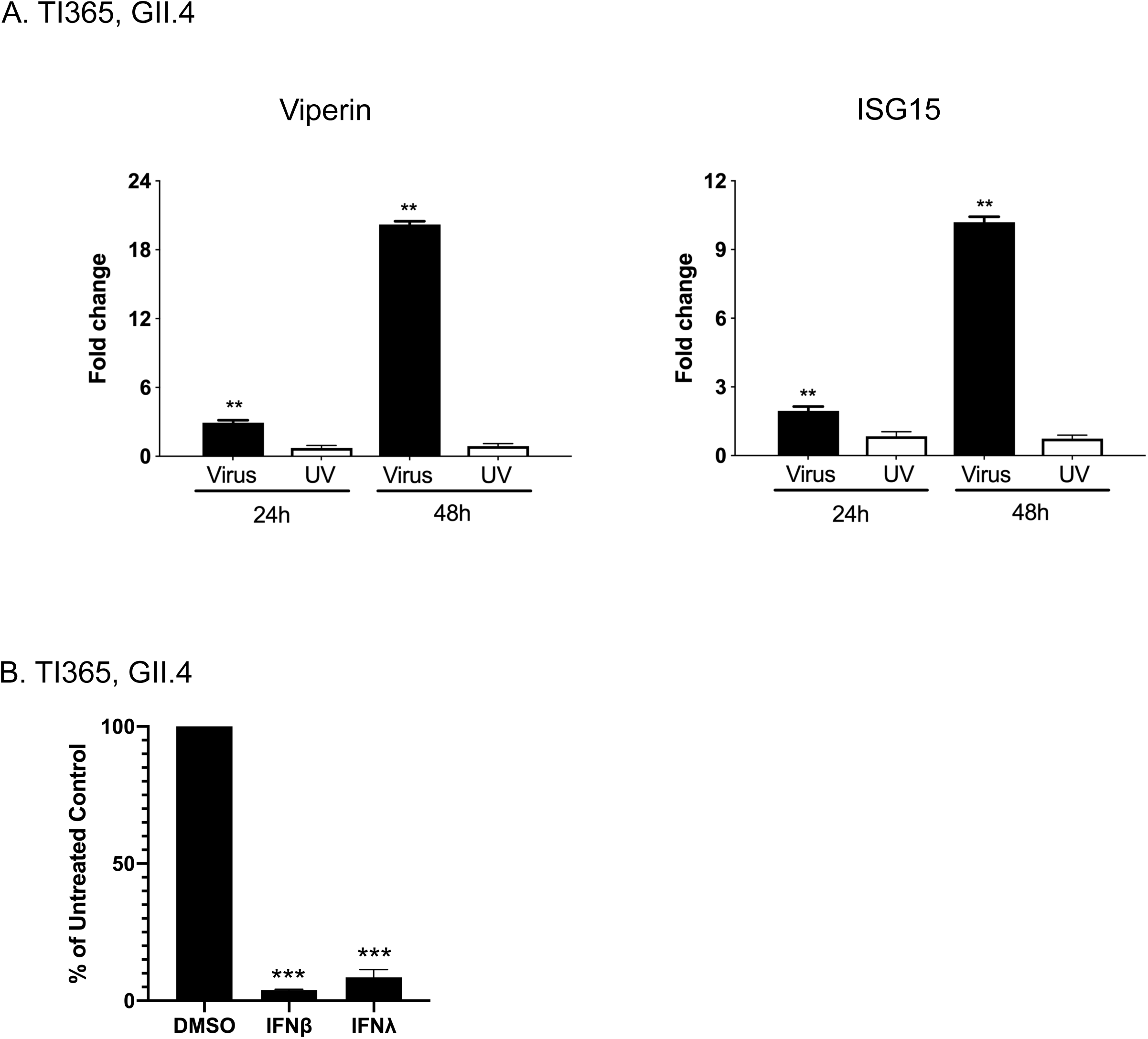
Human norovirus infection of intestinal epithelial cells induces interferon stimulated genes and is sensitive to Type I and III interferon. A, B) The levels of two interferon stimulated genes, viperin and ISG15, following norovirus infection of differentiated intestinal epithelial cells from the terminal ileum (TI365) was assessed at 24 and 48 hours post infection. Monolayers either GII.4 human norovirus or UV-inactivated GII.4. Relative gene expression was assessed by RT-qPCR and is shown as Log2 fold induction in comparison to the mock infected control. C) GII.4 Human norovirus infection of terminal ileum is sensitive to Type I (IFNβ) and Type III (IFNλ). Differentiated IEC monolayers were either mock treated or treated with recombinant IFN for 18 hours prior to infection with GII.4 HuNoV. Viral RNA levels at two days post infection were then quantified by RT-PCR and the increased in viral RNA expressed as a percentage of the untreated control.

To examine whether HuNoV replication in IECs was in fact sensitive to IFN and therefore by extension, likely restricted by the virus-induced response, we examined the effect of IFN pretreatment on the replication of HuNoV GII.4 in IECs. The addition of either IFNβ1 or IFNλ1/2 had an inhibitory effect on GII.4 HuNoV replication in IECs derived from terminal ileum organoids. This result confirms that HuNoV infection of IECs is sensitive to antiviral effects of type I (IFNβ1) or III (IFNλ1/2) IFN (Fig. 2B). We therefore hypothesized that the HuNoV-induced transcriptional responses may restrict HuNoV replication in IECs.

### Human norovirus replication in intestinal epithelial cells activates the IFN-induced JAK/STAT signalling pathway

The impact of norovirus infection on host gene expression in IECs and the magnitude of the IFN response during HuNoV infection, was examined by performing RNA-Seq analysis of infected IEC monolayers. IECs from two independent terminal ileum-derived organoid cultures were mock-infected, infected with a patient-derived GII.4 HuNoV strain, or with a UV-inactivated sample of the same inoculum. Two days post infection, total cellular RNA was extracted and processed for RNA-Seq analysis.

Robust infection of the cultures was evident from the increase in viral RNA levels over time with a 2753-fold and 498-fold increase in HuNoV RNA seen in the terminal ileum lines TI365 and TI006 respectively (Fig. 3A and 3B). As expected, very little increase in viral RNAs were observed in IEC monolayers inoculated with UV-inactivated stool filtrate (Fig. 3A and 3B). To identify genes differentially regulated in the response to productive HuNoV replication, pairwise gene comparisons from mock-infected, IEC monolayers infected with UV-inactivated HuNoV or HuNoV infected organoids were performed (Fig. 3C-3H). Three biological repeats of each condition was analysed by RNA-Seq as described in the methods. A total of 70 genes were found to be differentially regulated in GII.4 HuNoV IECs derived from organoid line TI365 when compared to the mock infected sample, with 69 increasing and 1 decreasing in their expression level (Fig. 3C, Table S3). Comparing the infected TI365 samples to the samples infected with UV-inactivated inoculum resulted in a slight increase in the number of differentially regulated genes; 76 in total with 73 increased and 3 decreased (Fig. 3E, Table S3). UV inactivation of the sample resulted in a near complete ablation of the transcriptional response when compared to the mock infected cells with very few reaching statistical significance, confirming that the transcriptional signature was virus-specific (Fig. 3G and 3H). In comparison, 162 genes were differentially regulated in GII.4 HuNoV-infected IECs derived from organoid line TI006 in comparison to mock infected cells; 9 decreased in expression and 153 increased (Fig. 3D, Table S3). The number of differentially regulated genes were reduced when compared to the IECs infected with UV-inactivated inoclum, 142 in total (Fig. 3F, Table S3). Similarly to our observations with GII.4 HuNoV infection of TI365, infection of TI1006 with the UV-inactivated inoculum resulted in a near complete loss of the transcriptional response.

**Figure 3.**
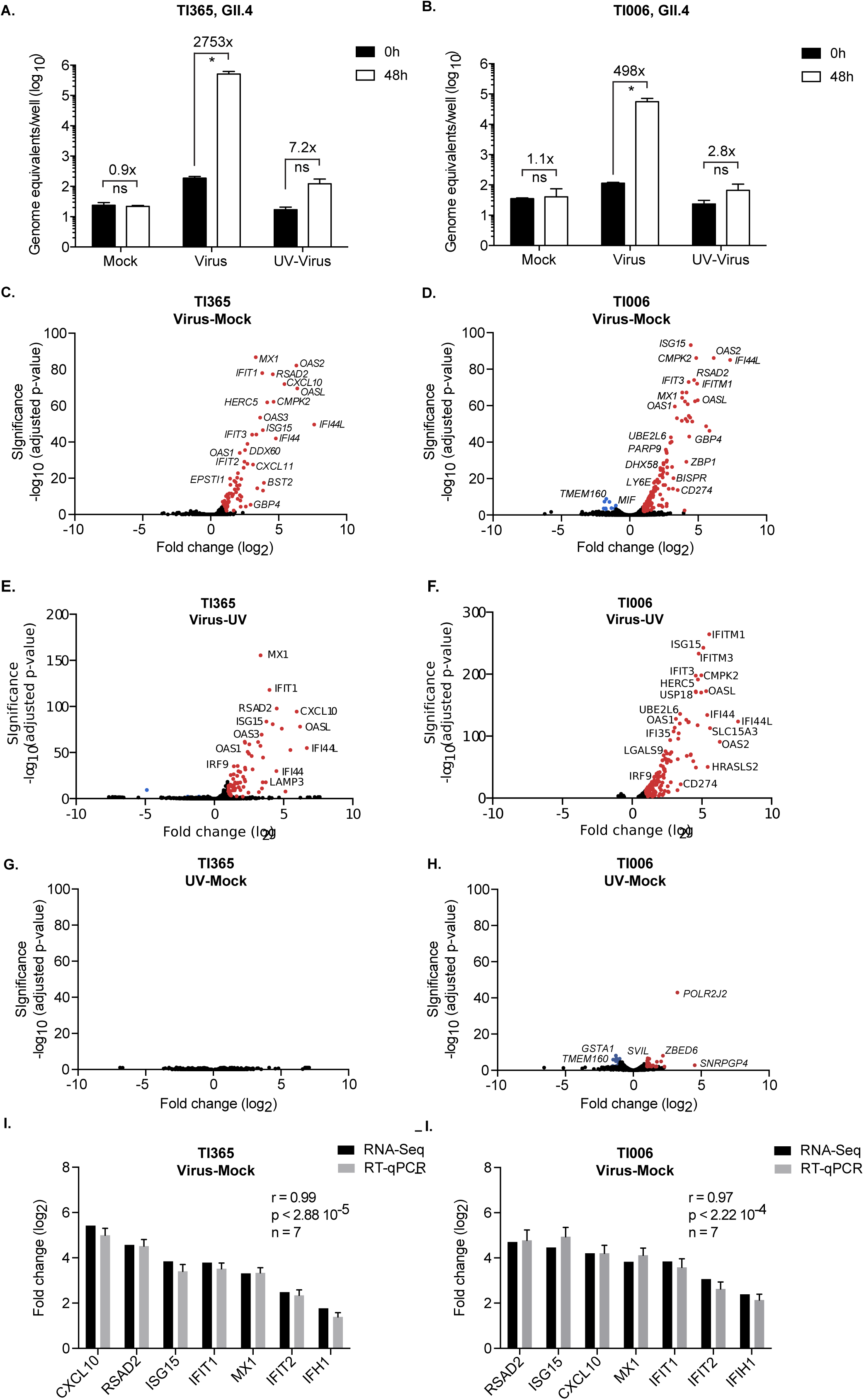
Norovirus infection of intestinal epithelial cells results in an interferon-induced transcriptomic response. A, B) IECs derived from two terminal ileum organoid lines (TI365 and TI006) were infected with a GII.4 HuNoV containing stool filtrate (Virus), the same stool filtrate that was UV-inactivated or mock infected and the levels of viral RNA quantified 48 hours post infection by RT-qPCR. Infections were performed in biological triplicate and quantified by RT-qPCR in technical duplicate. Error bars represent SEM. C-F) Volcano plots of differentially expressed genes from RNA-Seq analysis comparing gene expression in two different HuNoV infected organoids compared to mock (C, D) or UV-treated HuNoV infection (E, F). Significantly up- or down-regulated genes (FDR<0.01 and log2 fold change ≥ 1) are represented in red or blue, respectively. G-H) Comparison of expression changes of selected genes following HuNoV infection measured by RNA-Seq and RT-qPCR. Error bars represent the SD of one experiment performed in biological triplicate. The Pearson correlation coefficient (r), associated p-value (p) and the number of pairs analysed (n) are indicated on each chart.

In order to validate the results from the RNA-Seq analysis, we selected 7 differentially regulated genes and performed RT-qPCR on the same biological samples. We observed a strong correlation between RT-qPCR and RNA-Seq results confirming the accuracy of the expression data obtained by RNA-Seq (Fig. 3I and 3J).

Transcription factor enrichment analysis unambiguously identified STAT1 and STAT2 binding sites as highly enriched in promoter region of genes whose expression is significantly regulated following infection of human intestinal organoids (Fig. 4B). This strongly suggested that the JAK-STAT signaling pathway is activated following HuNoV infection. In agreement, gene ontology analysis highlighted a profound induction of type I interferon signaling in the transcriptomic response to HuNoV infection (Fig. 4C). In addition, Reactome pathway analysis carried out on the same set of genes further confirmed the involvement of interferon signaling in response to HuNoV infection (Fig. 4D).

**Figure 4.**
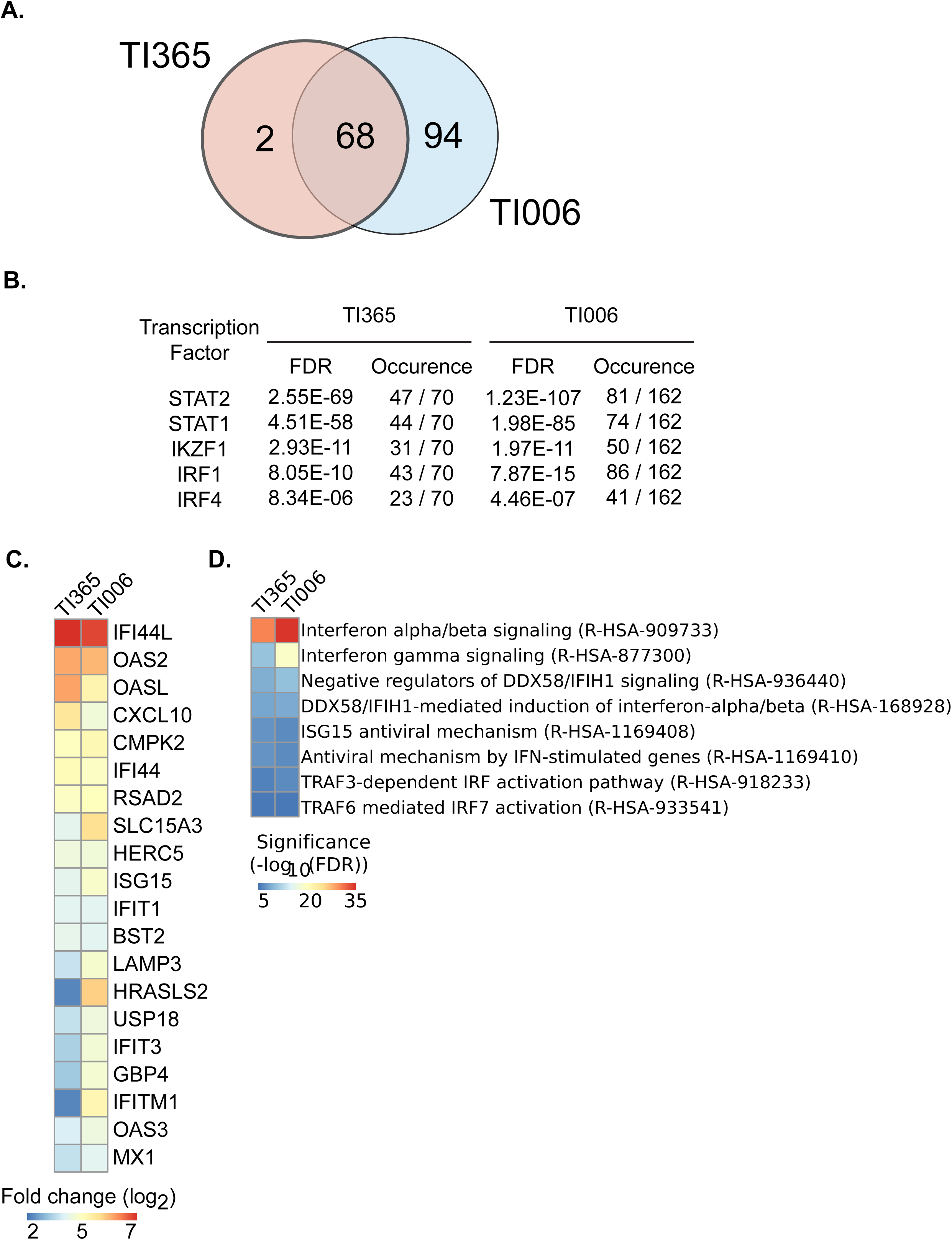
Human intestinal epithelial cells mount an interferon response to GII.4 human norovirus infection. A) Transcription factor enrichment analysis from differentially expressed genes of IECs from two terminal ileum-derived organoid lines (TI365 and TI006). Enriched transcription factors, number of occurrence among significantly regulated genes and significance are indicated for each organoid infection. B) Heat map showing the expression changes of the top 20 genes across two independent IEC infections. Genes are arranged by decreasing average enrichment fulfilling a false discovery rate (FDR) lower than 0.01. C) Heat map showing the most significant enriched Gene Ontology categories for biological processes inferred from significantly regulated genes across two independent organoid infections.

Comparing the genes differentially expressed in response to HuNoV infection with the Interferome database revealing that 94% (66) and 86% (140) of these genes were categorized as ISGs in organoid lines TI365 and TI006 respectively. Overall, these results demonstrated that HuNoV infection is readily sensed by IECs and that the IFN-induced JAK/STAT signalling pathway is likely activated during HuNoV infection and/or active replication.

### Genetic modification of intestinal organoids to ablate interferon induction or interferon signaling enhances HuNoV replication in IECs

To further examine the impact of IFN induction and the IFN signaling pathway on the restriction of HuNoV replication in IECs, we used lentiviral vectors to express viral innate immune antagonists to generate interferon deficient intestinal organoid lines. Lentiviral vectors were used to drive constitutive expression of either BVDV NPro or PIV5 V proteins, two well-characterized viral innate immune antagonists in a duodenum-derived organoid line (D196). In brief, the BVDV NPro protein originates from a non-cytopathic, persistent biotype of BVDV which effectively blocks IFN production by degrading IRF3, thereby preventing the activation of the innate immune system (40, 41). The PIV5 V protein instead targets IFN production as well as antiviral signaling by targeting STAT1, MDA5 and LGP2 for proteasomal degradation (42–45). To confirm that transduced proteins were functional, the expression levels of STAT1 and IRF3 were examined by western blotting. Whilst the STAT1 protein was present in the non-transduced control organoid line and BVDV NPro-expressing organoid lines, no STAT1 protein was observed in PIV5 V-expressing cells (Fig. 5A). The IRF3 protein was not detected in BVDV NPro-expressing cells, but was present in control and PIV5 V-transduced cells (Fig. 5A), confirming NPro protein functionality.

**Figure 5.**
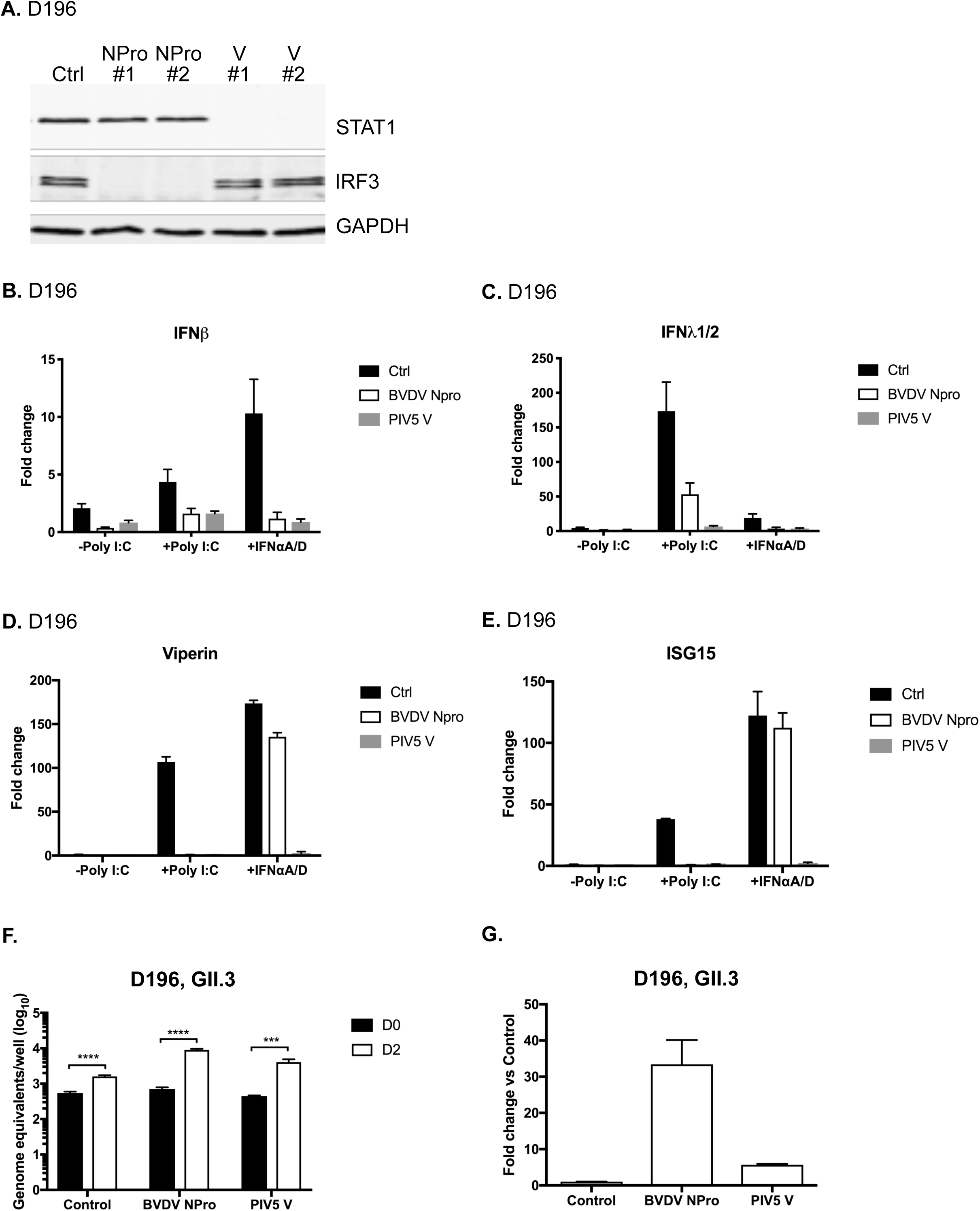
Genetically modified IFN-deficient organoids are more permissive for HuNoV replication. Duodenal intestinal organoids were modified by lentivirus-mediated transduction of the viral innate immune regulators, BVDV NPro and PIV5 V proteins. A) Two independent clones of transduced organoids were lysed and the expression of STAT1, IRF3 and GAPDH were examined by western blot to confirm the functionality of the BVDV NPro or V protein in intestinal organoids. B-E) To verify the inhibition of IFN production or IFN signaling, unmodified or modified intestinal epithelial cells were differentiated into monolayers and were transfected with poly I:C or treated with recombinant universal Type I interferon (IFNαA/D). The levels of IFNβ, IFNλ1/2, viperin and ISG15 were then quantitated by RT-qPCR. F, G) Replication of HuNoV GII.3 was examined in IECs derived from IFN-deficient organoids (D196). The levels of viral RNA obtained 48h post infection (D2) where compared to those obtained 2h post infection (D0). The levels of viral RNA replication seen in modified organoids were expressed relative to that seen in the control unmodified organoid line. All experiments were performed at least two independent times and results are expressed as mean ±SEM from duplicate samples analyzed in technical duplicate. Statistically significant values are represented as: *p ≤ 0.05, **p ≤ 0.01, ***p ≤ 0.001 and ****p ≤ 0.0001.

To verify the ability of NPro and V proteins to inhibit IFN induction and IFN signaling in the intestinal epithelium, IECs derived from stably transduced organoid lines were transfected with polyinosinic acid:polycytidylic acid [poly(I:C)] or treated with recombinant universal type I IFNαA/D a hybrid between human IFNα A and D. The levels of IFNβ, IFNλ1 and two representative ISGs, viperin and ISG15, then quantified by RT-qPCR. Following poly(I:C) transfection, elevated levels of IFNβ, IFNλ1, viperin and ISG15 mRNAs were observed in control IECs transduced with the empty vector as expected (Fig. 5B-5E). In comparison, the levels of IFNβ and IFNλ1 mRNAs induction was significantly lower in BVDV NPro- and PIV5 V-expressing cells, as were the mRNAs for viperin and ISG15 mRNAs (Fig. 5B-5E). Following treatment with type I IFN, IECs expressing the BVDV NPro or PIV5 V proteins showed significantly reduced IFNβ and IFNλ1 mRNA induction levels when compared with the control cells (Fig. 5B and 5C). Viperin and ISG15 mRNAs were not induced in IFN-treated PIV5 V-transduced IECs, confirming the impact of the V-protein in IFN signaling (Fig. 5D and 5E). These data further confirm that the IECs were IFN competent and that the BVDV NPro and PIV5 V proteins could efficiently block IFN induction and signaling in IECs.

The ability of HuNoV to infect IECs from the transduced organoid line was then investigated. Infection of the non-transduced D196 line with a GII.3 HuNoV strain resulted in only modest levels of virus replication; a ∼6-fold increase in viral RNA over 48 hours in the control non-transduced line was observed suggesting that the replication of this isolate in the D196 line was inefficient (Fig. 5F). However, suppression of the innate response by the expression of the N-Pro or V proteins stimulated GII.3 HuNoV replication in the D196 line; when comparing the yield of viral RNA from IECs derived from the non-transduced D196 line to those obtained from the transduced IECs, we observed that GII.3 HuNoV replication was increased by ∼33-fold and ∼6-fold in the NPro and V protein expressing D196-derived IECs respectively (Fig. 5G). Furthermore, we found that HuNoV GII.3 infection of the transduced lines did not induce ISG15, confirming the functionality of the transduced innate immune antagonists in HuNoV infected IECs (not shown). Similar results were obtained using a second transduced duodenal organoid line (results not shown). These results demonstrate that IECs produced from IFN-deficient organoids are more permissive for HuNoV and that the innate response limits HuNoV replication *in vitro*.

### Selective inhibition of Jak1/Jak2 enhances HuNoV replication in IECs

To further dissect the role of intestinal epithelial innate responses in the restriction of HuNoV infection, we investigated the effect of a specific Janus-associated kinase (Jak)1/Jak2 inhibitor on HuNoV replication in human IECs. Ruxolitinib (Rux), is an FDA approved drug of treatment for patients with dysregulated Jak signaling associated with myelofibrosis (46, 47) and for graft versus host disease (GvHD) (48). Rux has also been used to enhance growth of viruses that are sensitive to IFN (49). We first verified the ability of Rux to inhibit Type I and Type III IFN signaling following treatment of differentiated IEC monolayers derived from duodenal organoids with IFNβ or IFNλ1/2. Rux pretreatment was able to efficiently block the induction of viperin and ISG15 mRNAs following treatment with IFNβ or IFNλ1/2 (Fig.6A and 6B). We then examined the effect of Rux on HuNoV replication in IECs derived from the proximal duodenum and terminal ileum (Fig. 6C-6F). Differentiated IEC monolayers were inoculated with either a GII.3 or GII.4 HuNoV-positive stool filtrates. The inoculum was removed after two hours and cells were washed and maintained in bile acid (GCDCA)-containing media supplemented with either DMSO, Rux, or 2-*C*-methylcytidine (2-CMC). 2-CMC was included as a control as a well characterized inhibitor of HuNoV replication in replicon containing cells (26).

**Fig. 6.**
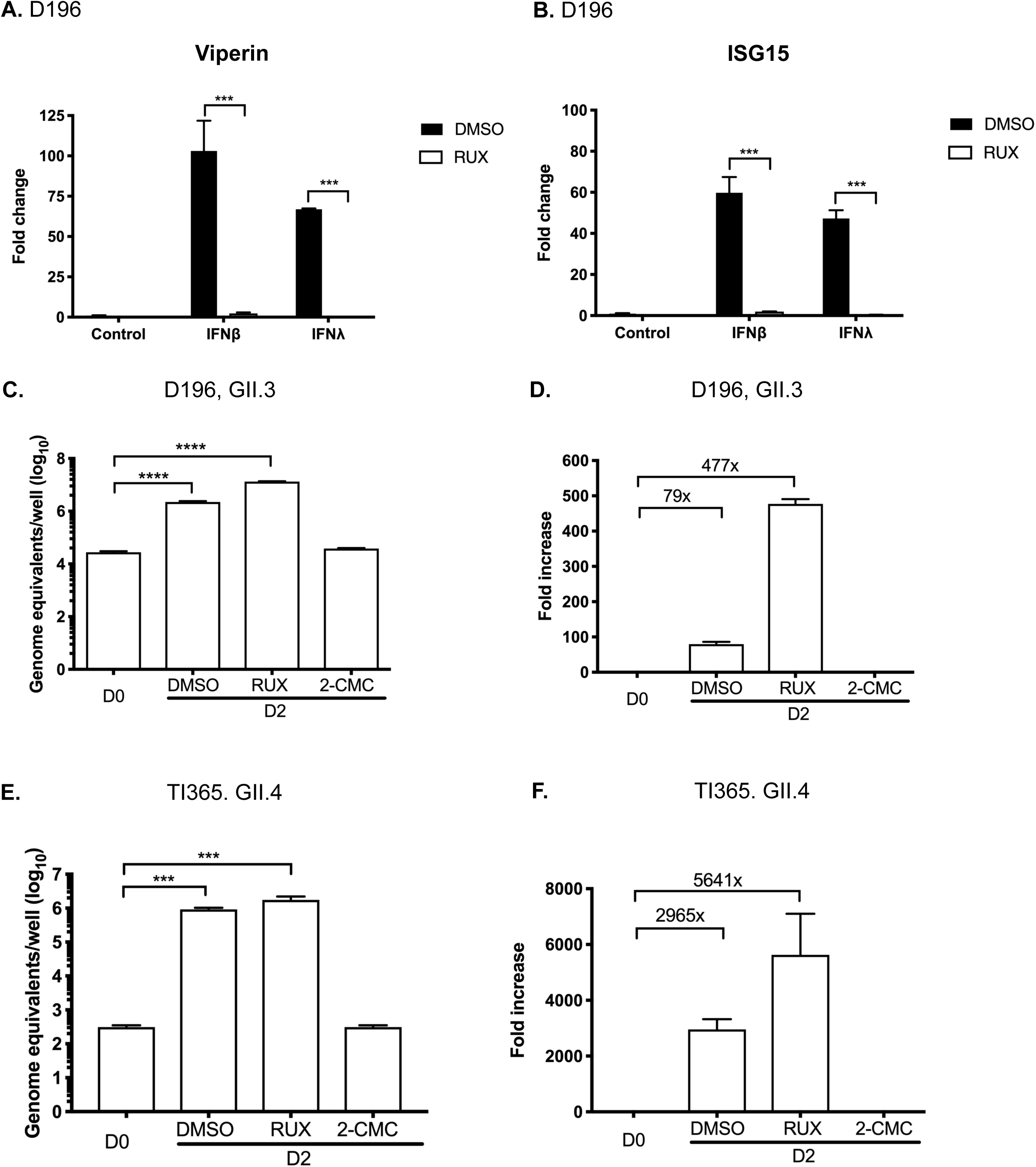
Inhibition of Jak1/Jak2 by Ruxolitinib (RUX) enhances HuNoV replication in intestinal epithelial cells. A,B) The ability of ruxolitinib (Rux) to inhibit type I and type III IFN signaling was examined following interferon (IFNβ or IFNλ1/2) pretreatments of intestinal epithelial cells derived from duodenal intestinal organoids (D196). C-F) To investigate the impact the role of JAK signaling in the restriction of HuNoV replication, intestinal epithelial cells were treated with DMSO, ruxolitinib (RUX) or 2-CMC (an inhibitor of HuNoV RNA-dependent RNA polymerase), and the impact of viral RNA synthesis was examined at 48h post inoculation (D2) by RT-qPCR. All experiments were performed at least three independent times and results are expressed as mean ±SEM from triplicate samples analysed in technical duplicate. Significant values are represented as:*p ≤ 0.05, **p ≤ 0.01, ***p ≤ 0.001 and ****p ≤ 0.0001

The impact of Rux treatment on HuNoV replication was assessed at 2d p.i. by qRT-PCR and confirmed that the inhibition of Jak stimulated HuNoV replication. In the absence of Rux, we observed a ∼79-fold and ∼2965-fold increased in HuNoV GII.3 and GII.4 viral RNA respectively in IECs derived from duodenal and terminal ileum organoids respectively (Fig. 6C-F). The inclusion of Rux in cultures following inoculation resulted in a significant improvement of HuNoV replication in all cases; GII.3 replication was increased to ∼477- and GII.4 replication increased to ∼5641-fold over a 48 hour period (Fig. 6C-F). In all cases, the addition of 2-CMC inhibited HuNoV replication, producing levels of viral RNA near identical to those observed at D0 p.i. (Fig. 6C-F). Rux stimulated the replication of a number of HuNoV isolates in IECs derived from a variety duodenum and terminal ileum organoid lines (Fig. S1). These results confirm that activation of the Jak1/Jak2 inhibits HuNoV replication and that pharmacological inhibition of this pathway increased HuNoV replication in culture.

### Inhibition of RNA polymerase II-dependent transcription increases HuNoV replication in IECs

To further assess the impact of *de novo* transcriptional responses on the restriction of HuNoV replication in IECs we examined the impact of Triptolide (TPL), a compound extracted from a traditional Chinese medicinal plant (Tripterygium wilfordii Hook F), on HuNoV replication in culture. TPL has potent immunosuppressant and anti-inflammatory activities, exhibiting a broad pharmacological effects against inflammation, fibrosis, cancer, viral infection, oxidative stress and osteoporosis (50, 51). TPL is known to have both antiproliferative and proapoptotic effects on a range of cancers (52, 53) and is reported to modulate the activity of many genes including those involved in apoptosis and NF-κB-mediated responses (51, 54). RNA polymerase II has recently been shown to be selectively targeted by TPL, although the mechanism by which TPL inhibits RNA polymerase II activity is yet to be fully elucidated (55). One of the known effects of TPL is the rapid depletion of short lived RNAs including transcription factors, cell cycle regulators and oncogenes (50, 55). Recent work has also confirmed that TPL treatment inhibits the innate response and stimulates vesicular stomatitis virus induced oncolysis (56).

To examine if the inhibition of transcription by TPL on the capacity of IECs to respond to IFN, the levels of viperin and ISG15 were assessed following treatment of cells with IFNβ or IFNλ. The mRNA levels of viperin was increased by 100- to 250-fold after treatment of IFNβ1 or IFNλ1/2 in DMSO-treated control IEC monolayers, but these increases were almost completely suppressed by the inclusion of TPL (Fig. 7A). A similar observation was made for ISG15, in that the inclusion of TPL potently inhibited the induction by interferon (Fig. 7B). These data confirm that TPL at concentrations that do not to affect overall cell viability is effective at suppressing the innate response in IECs.

**Fig. 7.**
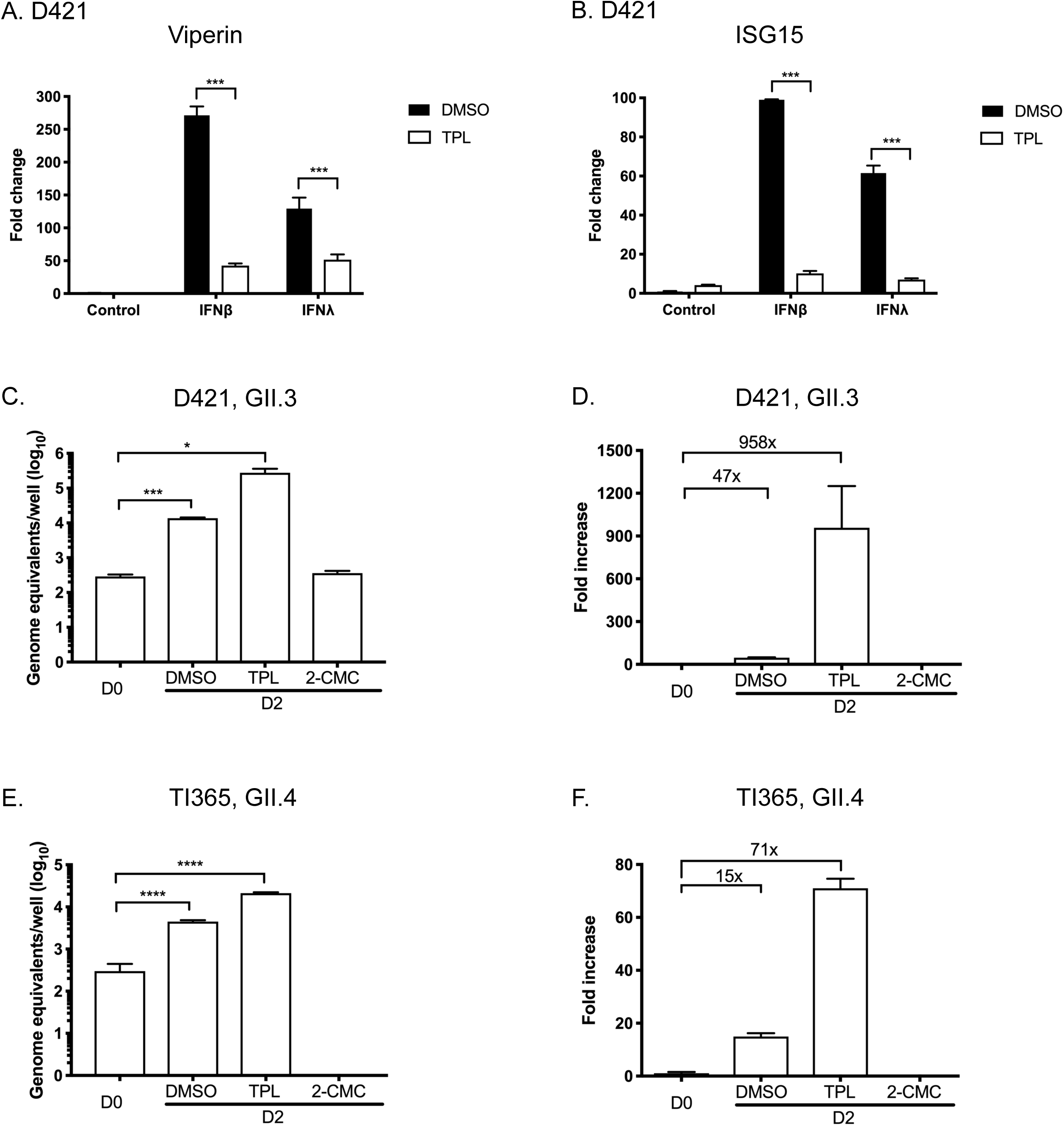
Inhibition of cellular transcription by triptolide (TPL), an RNAPII inhibitor, enhances HuNoV replication in intestinal epithelial cells. A,B) The ability of triptolide to inhibit type I and type III IFN signaling was examined following interferon pretreatment of intestinal epithelial cells derived from duodenal intestinal organoids (D421). C-F) To investigate the impact of restriction of transcription factors in HuNoV GII.3 and GII.4 replication, intestinal epithelial cells derived from duodenum (D421) and ileum (TI365) were treated with DMSO, triptolide (TPL) or 2-CMC (an inhibitor of HuNoV polymerase), and the impact of viral RNA synthesis was examined at 48h post inoculation (D2) by RT-qPCR. All experiments were performed at least three independent times and results are expressed as mean ± SEM from triplicate samples analysed in technical duplicate. Significant values are represented as:*p ≤ 0.05, **p ≤ 0.01, ***p ≤ 0.001 and ****p ≤ 0.0001

To examine the effect of TPL in HuNoV replication, differentiated monolayers generated from proximal duodenum and terminal ileum were inoculated with either GII.3 or GII.4 HuNoV-positive stool filtrates. After 2h, the inoculum was removed and cells were washed and maintained in GCDCA-containing differentiation media with either DMSO, TPL, or 2-*C*-methylcytidine (2-CMC) as a control. The addition of TPL resulted in enhanced HuNoV replication of GII.3 and GII.4 HuNoV strains (Fig. 7C-7F). These observations were consistent in IECs derived from both the duodenum (Fig.7C and 7D) and terminal ileum (Fig. 7E-7F). As expected, the HuNoV replication in the presence of 2-CMC was potently inhibited (Fig. 7C-7F). Enhancement of viral replication was also observed in another duodenum and terminal ileum organoid lines (Fig. S2). These results further confirm that inhibition of IFN-induced transcription increases HuNoV infection.

## Discussion

The efficient cultivation of HuNoV has remained a challenge since the initial identification of the prototype norovirus, Norwalk virus, in 1972 (57). Norovirus infection of the natural host species is very efficient, typically requiring <20 virus particles to produce a robust infection whereby >10^8^ viral RNA copies are shed per gram of stool within 24 hours (58, 59). Even in heterologous hosts (e.g. pigs) the HuNoV infectious dose has been estimated to be ∼ 2X 10^3^ viral RNA copies (60). Despite this, and despite enormous efforts, the ability to culture HuNoV efficiently has been a significant bottleneck in the study of HuNoV biology (57). Therefore, the ability to culture HuNoV has the potential to transform our understanding of many aspects of the norovirus life cycle, greatly enhance the capacity to develop therapeutics and allows the characterization of authentic viral neutralization titres following vaccination, rather than the current surrogate gold standard (21, 23). The net result of >40 years research has resulted in the establishment of two culture systems for HuNoV that use patient stool samples as the inoculum. The first such system relies on the replication of HuNoV within immortalized B-cells and requires the presence of enteric bacteria or soluble HGBA-like molecules from their surface (21, 22). Whilst, we have been able to reproduce the culture of HuNoV in immortalised and primary B-cells to varying degree of success (data not shown), we note that attempts by other labs have not universally been successful (21).

The recently developed HuNoV culture system IECs derived from intestinal organoids (23) while experimentally challenging, has been used in a number of subsequent studies to examine the impact of disinfectants (61) and the monoclonal antibodies (62, 63). This study set out to use organoid-based system to assess the cellular pathways that restrict HuNoV replication and to further refine the experimental conditions that allow optimal growth of HuNoV in culture. We found that HuNoV infection induces a robust innate response in IECs, in contrast to previous studies using transfection of purified HuNoV viral RNA into immortalized cells which concluded that the interferon response is unlikely to play a role (37). While the conclusions drawn in this previous study may be valid, it is likely that the inefficient replication seen using transfected RNA, where less than 0.1% of transfected cells contain active replicating viral RNA, reduce the sensitivity of the experimental system. This may be further confounded by unknown mutations that affect the robustness of the sensing pathways within immortalized cells. These previous observations, also contrast with our own findings that suggest that the ability of cells to respond to exogenous interferon negatively impacts on HuNoV replication (64, 65). This conclusion was based on the finding that the IFNλ receptor is epigenetically suppressed in an immortalized intestinal cell line which efficiently replicates a HuNoV GI replicon and that genetic ablation of IFNλ receptor expression enhances HuNoV replication in immortalized cells (65). We also recently described the generation of a robust culture system in zebrafish larvae, in which we also observed MX and RSAD2 (viperin) induction (39). Furthermore, it is well established that the interferon response is key to the control of MNV infection as mice lacking a competent innate response often succumb to lethal systemic MNV infections (66–68), demonstrating that the innate response is key to the restriction of norovirus infection to intestinal tissues in the mouse model (69). The development of the HuNoV organoid culture system provides the first opportunity to assess the impact of HuNoV infection on IECs, the first port of entry into the natural host.

Here we have seen that HuNoV infection of IECs induces an IFN-like transcriptional response by examining the replication of single HuNoV GII.4 isolate in IECs derived from two independent terminal ileum organoid lines from two different donors (Fig. 3). We chose the terminal ileum-derived organoids as our source of IECs as our data to date would suggest that GII.4 HuNoV replicates more efficiently in IECs derived from this gut segment whereas the GII.3 isolate replicated more efficiently in duodenal lines (Fig. 1D). Whether this difference was organoid line or viral strain specific, or suggests differing tropism is unknown, however this observation was consistent across several different duodenal or ileal organoid lines (data not shown).

Under the conditions used in the current study, the overall number of genes altered more than 2-fold in response to infection was relatively modest, 70 and 162 for TI365 and TI1006 respectively. We found that the transcriptional response induced in each organoid line was highly comparable, with a substantial overlap in the induced genes (Fig. 4). The use of UV inactivated inoculum allowed us to control for any non-specific effects of the other components of the filtered stool sample. Given the heterogeneity of any given stool sample, including this was essential to ensuring the observations were robust and represented alterations due to sensing of active viral replication intermediates. The rather modest number of genes induced, likely reflects the heterogenous nature of the IEC cultures and that not all cells in any given monolayer are permissive to infection. We estimate that ∼30% of cells were infected under the conditions used for the gene expression analysis which is similar to previous reports (23). The inclusion of Rux or TPL increased the overall number of infected cells in any given culture to ∼50% but even under the modified conditions, we have been unable to obtain higher levels of infection (data not shown). We hypothesize that obtaining higher levels of infection will likely require more uniform cultures, consisting primarily of enterocytes, the target cell for HuNoV (23).

The mechanism by which HuNoV is sensed by the infected cells is not currently known, however data from MNV suggests a clear role for Mda5-mediated sensing in the restriction of norovirus replication both in cell culture and *in vivo* (70). The sensing of MNV RNA occurs in a process that requires the HOIL1 component of the linear ubiquitin chain assembly complex (LUBAC) complex (71). Other components of the RNA sensing pathways have been implicated in the innate response to MNV including MAVS, IRF3 and IRF7 (70, 71) but the role they play in sensing of HuNoV RNA is unknown. In addition to targeting STAT1 for degradation (72), the PIV5 V protein is known to also inhibit the activity of Mda5 (44). Whilst not directly assessed, it is therefore likely that the stimulation is of HuNoV replication in the presence of the PIV5 V protein is a combined result of both of these activities. Further studies using gene edited organoid lines will be required to better define the relative contribution of each component in the sensing of HuNoV.

The most highly induced gene in response to HuNoV infection in both organoid lines was IFI44L, a novel tumour suppressor (73) previously show to have modest antiviral activity against HCV (74) and RSV (75, 76). IFI44L was also potently upregulated in IECS infected with human rotavirus (HRV) (77). Surprisingly, despite inducing a potent interferon response in IECs, HRV is not restricted by the endogenously produced IFN (77), an effect that has been hypothesized to be due to viral regulatory mechanisms that suppress the downstream activities of the induced genes. A number of the genes induced in response to HuNoV infection of IECs have previously been shown to have anti-viral activity against noroviruses. GBP4 and GBP1 were both induced following GII.4 infection of both organoid lines (Fig 3). The GBPs are interferon induced guanylate-binding proteins that are targeted to membranes of vacuoles that contain intracellular fungi or bacterial pathogens (78, 79), where they frequently result in the disruption of the pathogen-containing vacuoles (79). GBPs are targeted to the MNV replication complex in an interferon dependent manner that requires components of the autophagy pathway and exert their antiviral activity via an unknown mechanism (80). GBP2 was also identified as a norovirus restriction factor in a CRISPR based activation screen where it was found to have potent antiviral activity against two strains of MNV (81). Further studies will be required to determine if GBPs have similar antiviral effects during HuNoV infection.

The IFIT proteins IFIT1-3 were also significantly induced in response to HuNoV Infection of IECs (Fig 3, Table S1). The IFITs are a family of interferon stimulated RNA binding proteins that, at least in humans, are thought to inhibit the translation of foreign RNAs by binding to 5’ termini and preventing translation initiation (82, 83). In the context of norovirus infection, we have recently shown that the translation of norovirus VPg-linked RNA genome is not sensitive to IFIT1-mediated restriction (84), most likely due to the mechanism by the novel VPg-dependent manner with which norovirus RNA is translated (84). However, we did observe that IFIT1 in some way enhanced the IFN-mediated suppression of norovirus replication through an as yet undefined mechanism (84).

The development of the B-cell and organoid culture system have opened up the opportunity to dissect the molecular mechanisms of norovirus genome replication and to better understand host responses to infection. Others have observed that HuNoV replication in organoid derived IECs is highly variable (85) which agrees with our own experience during the course of the current study as we observed significant levels of week to week variation in infectious yield from the same organoid lines for any single strain of HuNoV (data not shown). We have also observed, as have others, that not all HuNoV strains appear to replicate efficiently in IECs derived from any single organoid line, which likely reflects the natural biology of HuNoV as individual susceptibility varies within any given population (85). What factors contribute to the relative permissiveness of any given organoid line to an isolate of HuNoV remains to be determined, but it is clear that Fut2 function appears essential for most HuNoV isolates as FUT2 negative lines were not permissive to the strains of viruses tested here (85, 86), data not shown). It is also possible that strains vary in the degree to which they induce and are sensitive to, the interferon response, as is common for other positive sense RNA viruses. Our data would suggest that irrespective of this, the replication of all isolates examined appear to be improved by treatment of cultures with Rux or TPL (Figs. 6 and 7; Figs. S2 and S3; and data not shown).

To our knowledge, our study represents the first demonstration that the genetic modification of human intestinal organoids can improve viral replication. The expression of BVDV NPro and PIV5 V proteins in cells has been widely used as a way to enhance virus replication in immortalised cells via the inactivation of aspects of the innate response (18, 87, 88). While the genetically modified organoids enhanced HuNoV replication by up to 30-fold in comparison to unmodified organoids we found that this varied between organoid lines examined (not shown). Surprisingly, we found that the process of differentiation, resulted in a significant increase in the basal levels of a number of ISGs (data not shown). Therefore the reason for variation in the enhancement is unknown but it may relate to the ability of any given organoid line to respond effectively and produce a rapid and effective innate response. The ability to readily generate gene edited human intestinal organoids while possible, is still very much in its infancy (89), therefore the ability to overexpress viral innate immune antagonists provides a more rapid way of generating intestinal organoids with specific defects in innate immune pathways. However, the simple inclusion of TPL or Rux appears to phenocopy the effect of overexpression of either NPro or V protein and can be readily applied to any organoid line. This low cost modification to culture conditions enhances the utility of the experimental system by improving the robustness of the replication.

The use of pharmacological inhibitors for the stimulation of viral infection has been described in many instances in immortalised cell lines (49, 56, 90), and more recently for viral infection of intestinal organoids (91). The mechanism of action of Rux is well defined as it specifically targets the JAK kinases (46). In contrast, the mechanism of action of TPL is less well defined but recent data suggests a direct mode of action on RNA polymerase II-mediated transcription (55). TPL has previously been shown to stimulate the replication of VSV by the inhibition of the interferon induced transcriptional responses (56). While TPL is not clinically used due to problems with water solubility, a water soluble pro-drug minnelide, has been trialled as an anti-cancer treatment for a number of cancers including pancreatic cancer (92).

Norovirus infection has now been widely accepted as a significant cause of morbidity and mortality in immunocompromised patients (13). In such cases, patients on immunosuppressive therapy following organ or stem cell replacement therapies, or those undergoing treatment for cancer, often suffer from infection lasting months to years (13, 93). Such infections have significant impact on the overall health of the affected patient, resulting in significant weight loss and a requirement for enhance nutritional support (94). Ruxolitinib, under the trade name Jakavi, is approved for the treatment of a range of diseases including splenomegaly in patients with myelofibrosis and has been shown to be effective in the treatment of chronic or acute (48, 95). Our data could suggest that the sustained administration of Rux in patients where chronic norovirus has been detected, may exacerbate the disease. We note however that during a study examining the effect of Rux on NK cell function in patients with STAT1 gain of function mutations, a single patient with chronic norovirus infection appeared to clear the infection following Rux treatment (96). The impact of Rux treatment on viral loads, and whether clearance was spontaneous, or due to improved NK cell function was not reported.

In summary, we have demonstrated HuNoV replication in IECs is restricted by the interferon response and that modulation of this response through either the genetic manipulation of intestinal organoids or the inclusion of pharmacological inhibitors, enhances HuNoV replication. Overall this work provides new insights into the cellular pathways and processes that control the replication of HuNoV, and provides improved conditions for the culture of HuNoV, enhancing the robustness of the HuNoV organoid culture system.

## Acknowledgments

This work was funded by research grants to IG: Wellcome Trust Ref: 207498/Z/17/Z IG is a Wellcome trust Senior Fellow. F.S. was funded by a Biotechnology and Biological Sciences Research Council (BBSRC) sLoLa grant (BB/K002465/1).

We thank Professor Steve Goodbourn (St. Georges Hospital, London) for the provision of the BVDV NPro and PIV5 V protein expression plasmids.

## Supplementary figure legends

**Figure S1: Ruxolitinib stimulates the replication of GII.4 in intestinal epithelial cells from a mucosa derived intestinal epithelial organoids (IEOs).** Intestinal organoids were generated from biopsies derived from the duodenum of four patients (Lines designated D353, D419, D421 and D428) or the terminal ileum of a single patient (TI006). Intestinal epithelial cell monolayers (IEC) were generated as described in the text and infected with either GII.3 or GII.4 human norovirus stool filtrate. The level of viral RNA levels were quantified at day 0 (D0) or 48 hours post infection (D2) by RT-qPCR and plotted as either genome equivalents per well (left hand column) or fold increase in viral RNA over 48 hours (right hand column). The impact of the inclusion of the Jak inhibitor (RUX) or the RNA polymerase inhibitor (2-CMC) on viral replication was examined by the addition of RUX or 2-CMC following the inoculation phase of the infection. All experiments were performed at least three independent times and results are expressed as mean ± SEM from triplicate samples analysed in technical duplicate. Significant values are represented as:*p ≤ 0.05, **p ≤ 0.01, ***p ≤ 0.001 and ****p ≤ 0.0001.

**Figure S2: Triptolide stimulates the replication of GII.3 or GII.4 in intestinal epithelial cells from a mucosa derived intestinal epithelial organoids (IEOs).** Intestinal organoids were generated from biopsies derived from the duodenum (Line D428) or the terminal ileum of a single patient (TI006). Intestinal epithelial cell monolayers (IEC) were generated as described in the text and infected with a GII.4 human norovirus stool filtrate. The level of viral RNA levels were quantified at day 0 (D0) or 48 hours post infection (D2) by RT-qPCR and plotted as either genome equivalents per well (left hand column) or fold increase in viral RNA over 48 hours (right hand column). The impact of the inclusion of the RNA polymerase II inhibitor (TPL) or the viral RNA polymerase inhibitor (2-CMC) on viral replication was examined by the addition of TPL or 2-CMC following the inoculation phase of the infection. All experiments were performed at least three independent times and results are expressed as mean ± SEM from triplicate samples analysed in technical duplicate. Significant values are represented as:*p ≤ 0.05, **p ≤ 0.01, ***p ≤ 0.001 and ****p ≤ 0.0001.

Supplementary Table 1. Supplier information of the composition of the organoid growth and differentiation media used for the culture of mucosal derived intestinal organoids in this study.

Supplementary Table 2. Supplier details of the interferons and inhibitors used in this study.

**Supplementary Table 3.** Summary of the differential gene expression analysis of GII.4 infected intestinal epithelial cells. RNA seq analysis was performed as described in the text. The data shown is the average of the data obtained from the two biological repeats. Differential gene expression analysis was performed by comparing the data obtained for infected vs mock infected IECs (Tabs A & C) or infected vs IECs infected with UV-inactivated inoculum.

